# Enemy release of introduced parasitoids does not affect their establishment or success

**DOI:** 10.1101/2024.11.05.621954

**Authors:** Miriam Kishinevsky, Mar Ferrer-Suay, Anthony R. Ives

## Abstract

The enemy release hypothesis, that the success of invading species is due to release from natural enemies that occur in their home range but not in the new range, can be an important explanation for successful invasions. Testing this hypothesis is difficult, however, because testing requires documented cases of not only successful, but also unsuccessful invasions. Therefore, observational data on after-the-fact establishment following unintentional introductions is insufficient, because they only include successful invasions. Here, we investigate the role of enemy release for parasitoid biological control agents and their hyperparasitoid natural enemies. We created a dataset of all aphid parasitoid wasp species introduced into North America, including data on their hyperparasitoid species in the native and new range. In total, information on 29 species of primary parasitoids and 54 species of hyperparasitoids was obtained, combining to create 259 parasitoid-hyperparasitoid species associations. Results show that introduced parasitoids experience partial enemy release, but the degree of enemy release does not affect the chance of successful establishment or successful control. This lack of effect of enemy release might be due to the broad geographical range of the hyperparasitoids (perhaps from unintentional introductions) which reduces the degree of enemy release. In addition, the life-history traits of the hyperparasitoid communities in the new range were different from the native range, with relatively more generalist hyperparasitoid species in the new range. These results show that introduced natural enemies can experience enemy release, but that this does not necessarily help to predict their successful establishment.

## Introduction

Invasive plants and animals can have major effects on the ecosystems they invade (Pimentel et al. 2000; Ricciardi and Atkinson 2004). One of the dominant hypotheses in invasion biology that explains successful invasions is the enemy release hypothesis (ERH) (Heger et al. 2024). This hypothesis states that if an invader species experiences a reduction in pressure from enemies in the new range, it will have an increased likelihood of successful establishment (Colautti et al. 2004; Prior and Hellmann 2015; Heger et al. 2024). This enemy release comes in the form of reduced pressure from predators, parasitoids, and/or pathogens attacking the invasive species in the new range (Torchin et al. 2003; Jeschke and Heger 2018). Underlying assumptions of ERH are that natural enemies play an important role in regulating the population of the invader in its home range, and that there is a loss of pressure from enemies in the new range (Roy et al. 2011; Prior et al. 2015). With this reduction in top-down regulation, invasive species can better succeed in the invaded range (Torchin et al. 2003). Most studies investigating enemy release focus on invasive plants, with many of them finding evidence of enemy release (Connor et al. 1980; Keane and Crawley 2002; Prior and Hellmann 2015). In insects, ERH was studied in an invasive fire ant species (Yang et al. 2010), mosquitos (Aliabadi and Juliano 2002) ladybeetle species (Koyama and Majerus 2008), gall wasps (Schönrogge et al. 2000; Prior and Hellmann 2013), and invasive herbivorous insects (Cornell and Hawkins 1993) (Table 12–a1 in Prior and Hellmann 2015).

The presence and consequence of enemy release depend on characteristics of both the invader and the range it is invading. In a comparative study of parasitism of invasive herbivorous insects, (Cornell and Hawkins 1993) showed that the number of parasitoids in the new range attacking the invasive herbivores was correlated with the number attacking the species in their native range. The authors called this characteristic ‘vulnerability-to-parasitism’ that could be explained by both the intrinsic characteristics of the host and by extrinsic factors like the taxonomic match of the parasitoid species pool to the host, particularly if hosts have closely related species in the new range that are already attacked by native parasitoids. It is also possible for enemies in the new range to include more generalists that attack a broader range of hosts, including the invading species (Keane and Crawley 2002; Roy et al. 2011).

Because enemy release depends on characteristics of the invaded range, enemy release is likely to change through time, but studies of enemy release through time are rare (Barney 2006; Schulz et al. 2019). Enemy release may be temporary, decreasing with time since invasion as more locally occurring enemy species transfer to attack the invader (Schönrogge et al. 2000; Aliabadi and Juliano 2002; Torchin et al. 2003). Thus, not only does enemy release depend on the characteristics of both the invader and the new range, these characteristics may change and lead to differing short- and long-term importance of enemy release (Jeschke and Heger 2018).

Although ERH is well recognized, a lot of the evidence for ERH is observational, with studies finding that successful invasive species have fewer enemies in the new range and concluding that this benefited the invaders (Prior and Hellmann 2015). A problem with making this inference from observational data is the fact that species invasions are mostly unintentional, and what determines successful versus failed invasions is often hard to diagnose. In contrast to unintentional introductions, importation (classical) biological control provides cases of intentional introductions of alien species that can give information about ERH (Taylor and Elton 1959; Roy et al. 2011; Heimpel and Mills 2017).

### Parasitoids are one of the main taxonomic groups used for importation biological control

In part this is because they are on average more specialized than predatory arthropods and therefore are less likely to have unintended consequences in the new range (Waage and Hassell 1982). One of the potential disadvantages of parasitoids as biological control agents, however, is that they are often attacked by hyperparasitoids, secondary parasitoids that parasitize primary parasitoids (Fiske 1910; Boivin and Brodeur 2006). Hyperparasitoids are ubiquitous in insect terrestrial food webs across a wide range of environments (Tougeron and Tena 2019; Poelman et al. 2022). In some cases they have been found to interfere with biological control efforts (examples in Table 1 of Stiling 1993), but in other cases biological control was successful despite hyperparasitoid activity (Hougardy and Mills 2009; Broadley et al. 2018). For example, two species of ichneumonid parasitoids were introduced into North America to control the alfalfa weevil, *Hypera postica* (Gyllenhal, 1813). One of the species, *Bathyplectes curculionis* (Thomson, 1887), had 20 species of hyperparasitoids attacking it. The other species, *B. anurus* (Thomson, 1887), had lower numbers of hyperparasitoids due to its different overwintering strategy, and *B. anurus* was the more effective biological control agent (Day 1981; Radcliffe and Flanders 1998). In other examples, primary parasitoids were introduced and were able to provide successful biological control even when hyperparasitism rates were very high. Hyperparasitism reached 90% for the introduced parasitoids of the cassava mealybug and the mango mealybug in some areas of Togo (Fischer 1991), and up to 100% in some periods of the growing season for the introduced parasitoid of the walnut aphid in California, USA (Bosch et al. 1979); nonetheless, in both cases parasitoids were able to provide effective biological control. Despite example cases of hyperparasitoids both hampering and having no effect on importation biological control, there is still no answer to the general question of how frequently hyperparasitoids interfere with successful importation biological control (Schulz et al. 2019).

**Table 1.**
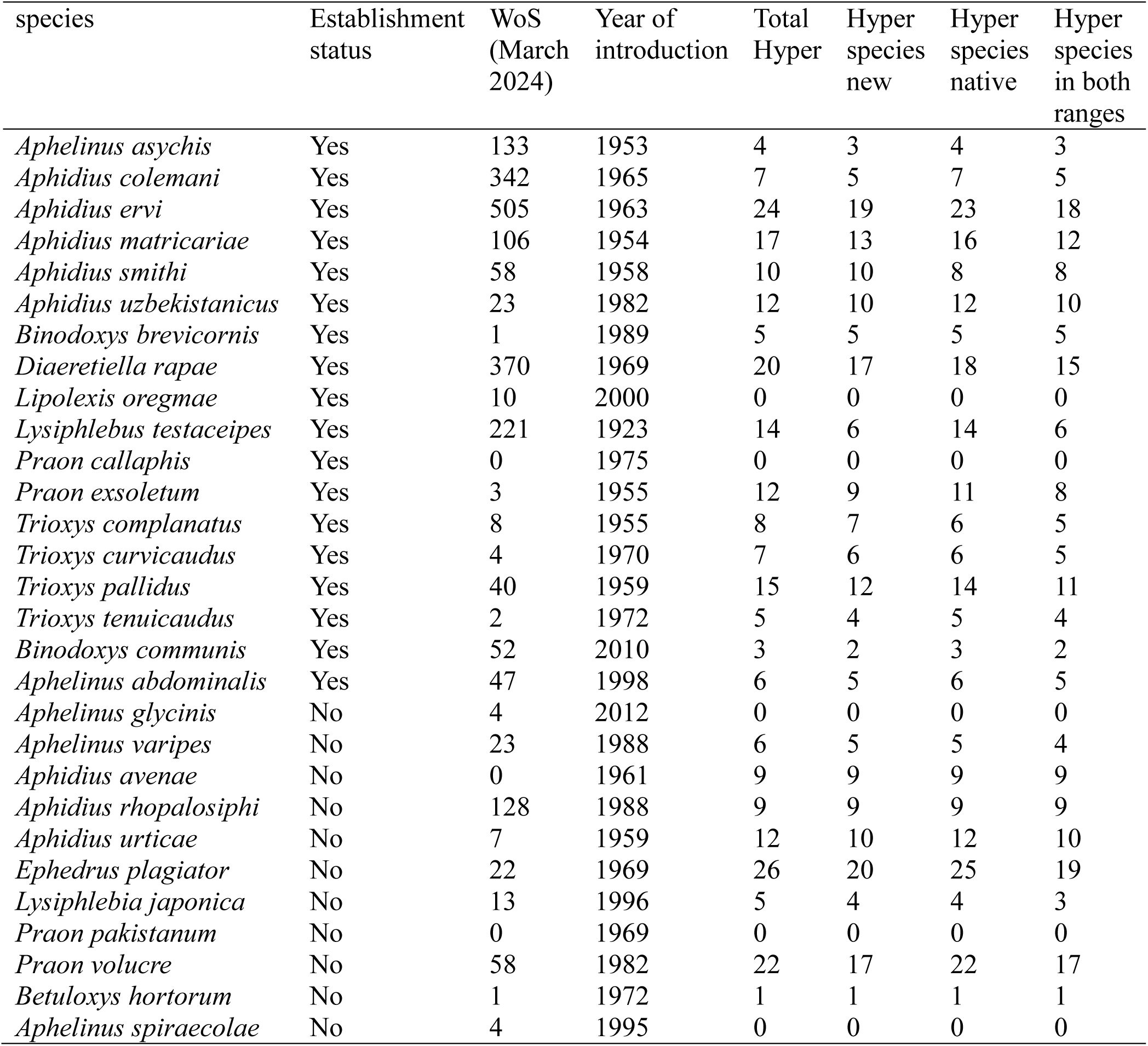
Species of primary parasitoids of aphids introduced into North America including Hawaii. The columns give the number of articles for each primary parasitoid species in the Web of Science (WoS) at the time searched, the year the primary parasitoid was introduced for biological control, the total number of hyperparasitoids (Hyper) and the numbers in each range, and the number of hyperparasitoids occurring in both ranges.

Here, we investigate whether parasitoids used for importation biological control experience enemy release in the invaded range by conducting a biogeographical comparison of ERH (Prior and Hellmann 2015). We focused on primary parasitoids that were released into North America for biological control of aphid pests; as a group, these parasitoids have a large and taxonomically diversity collection of hyperparasitoids (Brodeur 2000). We asked whether aphid primary parasitoids encounter fewer hyperparasitoid species after introduction into North America compared to their native range, and whether having fewer hyperparasitoids is associated with their ability both to establish and to provide biological control. We tested whether enemy release is at play for these parasitoid invaders, with the expectation that hyperparasitoids which are present in both the native and new range of the primary parasitoid would not allow for enemy release. In addition, we asked whether enemy release changes through time after introduction.

Finally, we compared the traits of hyperparasitoids (host range and whether hyperparasitoids attack larvae or pupae) which the invasive parasitoids encounter in the new range to the hyperparasitoids in the native range. General summaries of importation biological control have been published previously (Cock et al. 2016; Van Driesche et al. 2020; Jarrett and Szűcs 2022), but these summaries have not focused on hyperparasitoids and ERH.

## Methods

### Primary parasitoid data

Hymenopteran primary parasitoids of aphids (Aphididae) belong to two groups: all species in the braconid subfamily Aphidiinae (Braconidae) and some species in the chalcid family Aphelinidae (genus *Aphelinus* and more) (Sullivan and Völkl 1999; Sullivan 2009). We obtained data from the BIOCAT 2010.4 dataset, courtesy of Matthew Cock (Greathead and Greathead 1992; Cock et al. 2016), the ‘Catalog of Species Introduced into Canada, Mexico, the USA, or the USA Overseas Territories for Classical Biological Control of Arthropods, 1985 to 2018’ (Van Driesche et al. 2020), and Arredondo-Bernal and Rodríguez-Vélez (2020). All data on aphid primary parasitoids released to North America were used. Data from these three databases resulted in a total of 29 primary parasitoid species released between the years 1923 and 2012. Data on the native range of primary parasitoids were obtained from the same databases.

The primary parasitoids were introduced from Africa, Asia, and Europe; we did not consider species released into continental USA from other regions in continental USA (‘intracontinental introductions’ Prior and Hellmann 2013). Species that originated in non-continental USA and were introduced into the mainland contiguous states were also included (1 species from Guam) and vice versa (1 species introduced into Hawaii). For releases into Mexico from the USA after having been previously introduced into the USA, only the first introduction was considered (see Arredondo-Bernal and Rodríguez-Vélez 2020). Some species were released more than once, and some were released in different locations in North America (for example, different states in continental USA). The species *Aphelinus atriplicis* (Walker, 1839) was excluded because it was misidentified as *Aphelinus albipodus* (Hayat & Fatima, 1992) in most of the literature in the USA (Van Driesche et al. 2020), and therefore records of hyperparasitoid associations are questionable. *Praon gallicum* (Starý, 1971) was also excluded because its origin, and whether it is native or introduced into North America, are unclear (Acheampong et al. 2012).

### Hyperparasitoid data

Hymenopteran hyperparasitoids of aphid parasitoids are much more taxonomically diverse than aphid primary parasitoids, and we focused only on those with host records including aphid parasitoids that were introduced into North America (Canada, the USA and Mexico) for biological control. Aphid hyperparasitoids can be divided into two main groups. The first are referred as true hyperparasitoids (Gagic et al. 2012): they are koinobiont endo-hyperparasitoids that attack the primary parasitoid larva while the aphid is still alive. These are members of the subfamily Charipinae (Cynipoidea: Figitidae), specifically the genera *Alloxysta*, *Phaenoglyphis*, and *Lytoxysta*. The second group are sometimes called mummy parasitoids, and they are idiobiont ecto-hyperparasitoids attacking the prepupal and pupal stages of the primary parasitoid when the aphid is already mummified (dead). Two groups display this life strategy: Megaspilidae (Ceraphronoidea), specifically *Dendrocerus* and *Aphanogmus*, and Pteromalidae (Chalcidoidea), specifically *Asaphes*, *Pachneuron*, *Coruna*, and *Euneura* (Sullivan and Völkl 1999; Sullivan 2009). There are also members of the genus *Syrphophagus* (Chalcidoidea: Encyrtidae) that are endo-ecto koinobiont parasitoids able to attack primary parasitoid hosts both inside the living aphid and inside the dead mummified aphid, and in both scenarios the egg is laid inside the host and the larvae first feeds internally and later develops externally (Buitenhuis et al. 2004).

Although species from the genus *Syrphophagus* can do both, they mainly attack the mummy stage, and offspring fitness is higher when mummies are attacked (Buitenhuis et al. 2004); thus, species of *Syrphophagus* were classified as mummy hyperparasitoids. Very limited data exist about the aphid hyperparasitoid *Aprostocetus populi* (Kurdujmov 1913) (Chalcidoidea: Eulophidae) (Hagen and Van Den Bosch 1968; Yegorenkova et al. 2007), and its developmental mode is unclear. Another group for which records of aphid hyperparasitoids exist are *Marietta* species (Chalcidoidea: Aphididae) (Summy et al. 1979) which are also ectoparasitoids (Kfir and Rosen 1981), but there are only a few records of this association in the literature. There are groups of aphid hyperparasitoids excluded above, because they parasitize primary parasitoids other than our focal 29 species. Unless stated otherwise, we will refer to all of these groups as hyperparasitoids.

To gathered data on hyperparasitoid species, we searched Web of Science (data search ended August 1^st^ 2024) for the name of every primary parasitoid species (29 species) with double quotation marks, both for the full species name (e.g., *Aphidius ervi*, *A. ervi*) together with the word “hyperparasitoid”. If there were no results, the quotation marks were dropped. All retrieved documents were searched for hyperparasitoid associates of the 29 primary parasitoid species. In addition, for the primary parasitoids from the Chalcidoidea superfamily, the Universal Chalcidoidea Database (Noyes 2022) was searched for associated hyperparasitoid species and confirmed with the literature citing the association. Keys to species (which often have information about hosts) for the hyperparasitoid groups were searched for each of the primary 29 species. From these keys the range of the hyperparasitoid species was collected, and the possible association with each primary parasitoid hosts was categorized as: (i) hyperparasitoid only occurs in the native range of the primary parasitoid; (ii) hyperparasitoid only occurs in the new range of the primary parasitoid; (iii) hyperparasitoid occurs in both native and new ranges of the primary parasitoid. Details about the sub-continental range were disregarded, and in the case that the new range was continental USA, Canada or Mexico, hyperparasitoids that were present anywhere in that range were counted as “present in new range”. This implicitly assumes that these hyperparasitoid species occur throughout North America. While this is unlikely to be true for all hyperparasitoid species, hyperparasitoids are highly mobile, and in the absence of good data from this understudied group, we made this pragmatic assumption. Most of the data on associations and ranges came from keys, catalogues, and databases (Fergusson 1981; Gibson and Vikberg 1998; Ferrer-Suay et al. 2012, 2017, 2023; Noyes 2022). For all hyperparasitoid species, data on whether they attack the host inside the living aphid or attack the host inside the mummy were collected. For some primary and hyperparasitoid species, names were changed or synonymized over the years since identification. For these cases, we included the old names in our data search. Data from studies were excluded if more than one primary parasitoid was collected and the association of the hyperparasitoids was unclear between the species of the primary hosts.

One of our questions was whether hyperparasitoids with a greater host range are also geographically more widespread (occur in both native and new ranges). This question can be asked more broadly for other hyperparasitoid groups. We selected an entire subfamily of hyperparasitoids, the Charipinae (mostly species of *Alloxysta*), because this is a well-studied group by Ferrer-Suay et al. (2012). We assembled data for this hyperparasitoid group on the number of host species they attack and the number of biogeographical realms they occupy (Afrotropical, Australasian, Nearctic, Neotropical, Indomalayan, and Palearctic). This allows a comparison with the hyperparasitoids that attack aphid parasitoids as to whether hyperparasitoids that attack more species are more likely to occur in both native and new ranges.

The phylogeny of the parasitoid species can be associated with the interactions they have with hyperparasitoids and potentially their establishment and success as control agents.

Therefore, we constructed a phylogeny for the 29 primary parasitoid introduced species (according to Kambhampati et al. 2000; Stanisavljević et al. 2006; Heraty et al. 2007; Derocles et al. 2012; Hopper et al. 2012; Gokhman et al. 2017; Žikić et al. 2017; Mitrovski-Bogdanović et al. 2021; Čkrkić et al. 2021) and used it in the statistical analyses to account for phylogenetic correlations among species. Two other factors that could potentially affect the number of associations between parasitoids and hyperparasitoids are the amount of research performed on each species and the years since the species was described, assuming that more research and time since discovery could both increase the number of associations that exist on record in the literature. These two factors represent sampling bias, rather than the real number of associations. We used numbers of articles found in Web of Science (WoS) when searching the full name of the species with quotation marks, and the year the primary species was described, as independent variables (covariates) to factor out these sources of bias.

### Statistical analysis

The central question of this study is whether parasitoids that were introduced into a new range experience enemy release and whether enemy release effected establishment. We calculated the ‘degree of enemy release’ as the difference between the number of associations with hyperparasitoids in the native and new range divided by the square root of the sum of hyperparasitoids in both ranges; dividing by the square root of the total standardizes the variance in the response variable. The degree of enemy release was analyzed using a phylogenetic regression, with articles from WoS and year the species was described as covariates. This analysis is similar to a phylogenetic paired t-test with two covariates. For the analysis, we used a phylogenetic generalized linear mixed model (PGLMM) using the function pglmm_compare in the package ‘phyr’ (Li et al. 2020).

To test if establishment (yes/no) of introduced species was affected by enemy release, we used a binomial PGLMM (Li et al. 2020). For this test, there are no data on new associations for non-established species, and therefore we calculated the degree of enemy release excluding hyperparasitoid species that were only present in the new range for both non-established and established species. Thus, this analysis assumes that the release comes from associations that are present in the home range but not in the new range. Since it is also possible that hyperparasitoids of the released species occur in North America but do not occur at the site of release, we also did a similar analysis but with total enemy release, excluding all hyperparasitoids in the new range and therefore testing if the number of associations in the home range affects establishment. The number of articles in WoS for each species and year the species was described were included as independent variables. We similarly tested the effect of the degree of enemy release on the biological control success (yes/no); in this test, we calculated the degree of enemy release including hyperparasitoids that occur in both native and new ranges.

To test for a relationship between the numbers of hyperparasitoid species in the new range and in the native range associated with the established primary parasitoids, we used cor_phylo() that allows correlations among variables with phylogenetic signal (Li et al. 2020). We performed a similar analysis for the correlation between the degree of enemy release and the time since primary parasitoids were released to see if enemy release diminished with time.

To test whether the life histories of the hyperparasitoid species that attack the introduced primary parasitoids was different between the native and new range, we compared the arcsine-square root transformed proportion of mummy hyperparasitoids of established species in the native versus new range using a phylogenetic paired t-test with phyl.pairedttest in phytools (Revell 2012). Using the same method, we tested whether the hyperparasitoid species attacking in the new range included more generalist species; as a measure of whether hyperparasitoids were generalists, we used the number of hosts in the collected database for each hyperparasitoid species and calculated the mean host number of all hyperparasitoid species associated with each primary species.

All analyses were performed in R version 4.1.2 (R Core Team 2013).

## Results

In total, we found 54 species of hyperparasitoids that attack at least one of the 29 primary parasitoid species released to control aphids in North America. This resulted in a total of 259 associations, with the range of 0-26 hyperparasitoid species associated with each primary parasitoid (Table 1). For five of the primary species, no hyperparasitoid associations were found in the literature. The majority of the associations (74%) were the same in the native range and the new range due to the cosmopolitan nature of the generalist hyperparasitoids (Fig. 1a). Almost half of the associations (48%) included one of nine generalist hyperparasitoid species, with each of these hyperparasitoid species having more than 10 associations. Most hyperparasitoids had fewer associations, with 25 of them having only one. A positive relationship was found between the number of primary parasitoids which a hyperparasitoid attacks and the number of associations that occur in both native and new ranges (Fig. 1a). This relationship is similar to the relationship between the number of parasitoids attacked and the geographic ranges of all species of the hyperparasitoid subfamily Charipinae (mostly species of *Alloxysta* and *Phaenoglyphis*) (Fig. 1b).

**Figure 1.**
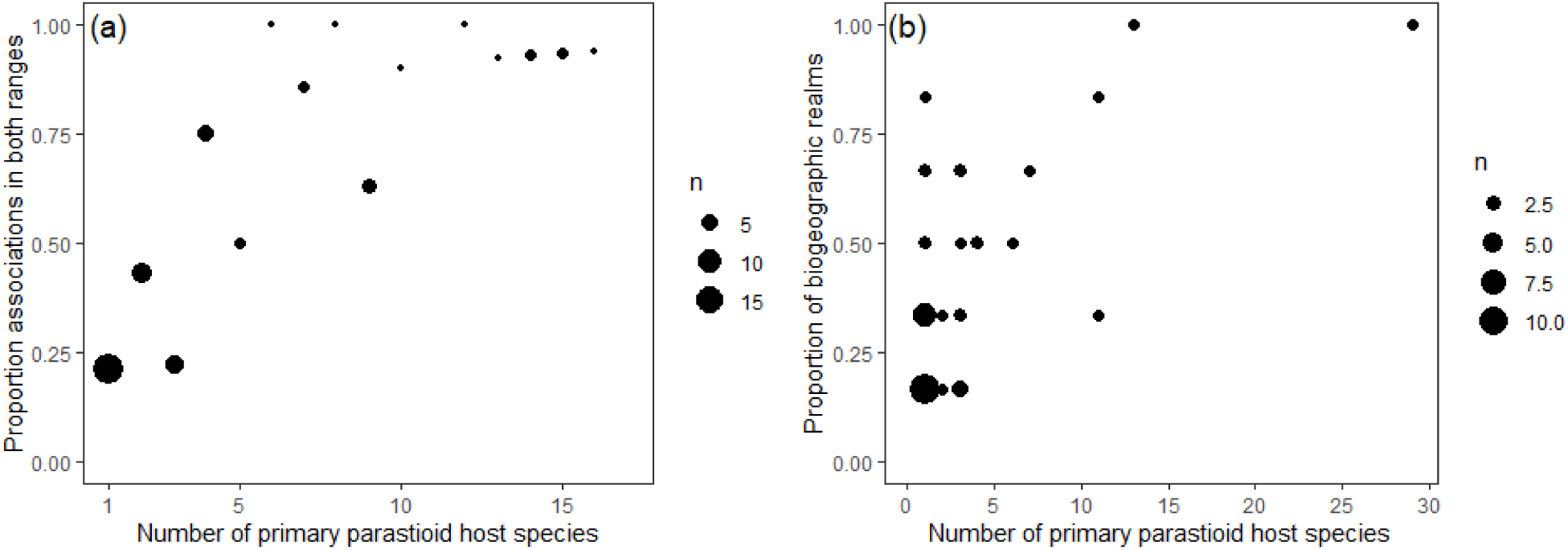
(a) For hyperparasitoid species attacking the 29 aphid parasitoids, the proportion of hosts occurring in both native and new ranges versus the number of parasitoids the hyperparasitoids attack. (b) For species of hyperparasitoid Charipinae, the proportion of biogeographic realms in which they have been recorded (Afrotropical, Australasian, Nearctic, Neotropical, Indomalayan, and Palearctic) versus the number of parasitoids they parasitize (data from Ferrer-Suay *et al*., 2012 and Ferrer-Suay *et al*., 2023). The size of the dots is proportional to the number of hyperparasitoid species (*n*).

Of the 29 parasitoids introduced for biological control, only 18 became established. Of the 18 established species, 14 had some enemy release, with 33 hyperparasitoid species and 51 associations present only in the native range of the primary parasitoid. The average number of hyperparasitoids associated with the established primary parasitoids was lower in the new range (7.4) than in the native range (8.8) (PGLMM, Z = 2.96, *p* = 0.003, with a phylogenetic random effect of 0.23) (Fig. 2a). Out of these 33 hyperparasitoid species, 18 belong to the subfamily Charipinae. Despite the evidence for enemy release in the established group, the degree of enemy release did not significantly affect establishment in the new range (Table 2, PGLMM, Z = 0.82, *p* = 0.4, with a phylogenetic random effect of 0.53); this analysis excludes hyperparasitoids that only occur in the new range because there are no data on new associations for non-established species. In the analysis assuming total enemy release (using only the number of hyperparasitoids in the native range), establishment was also not affected by the number of associations in the home range (Table 2, PGLMM, Z = 0.24, *p* = 0.8, with a phylogenetic random effect of 0.34).

**Figure 2.**
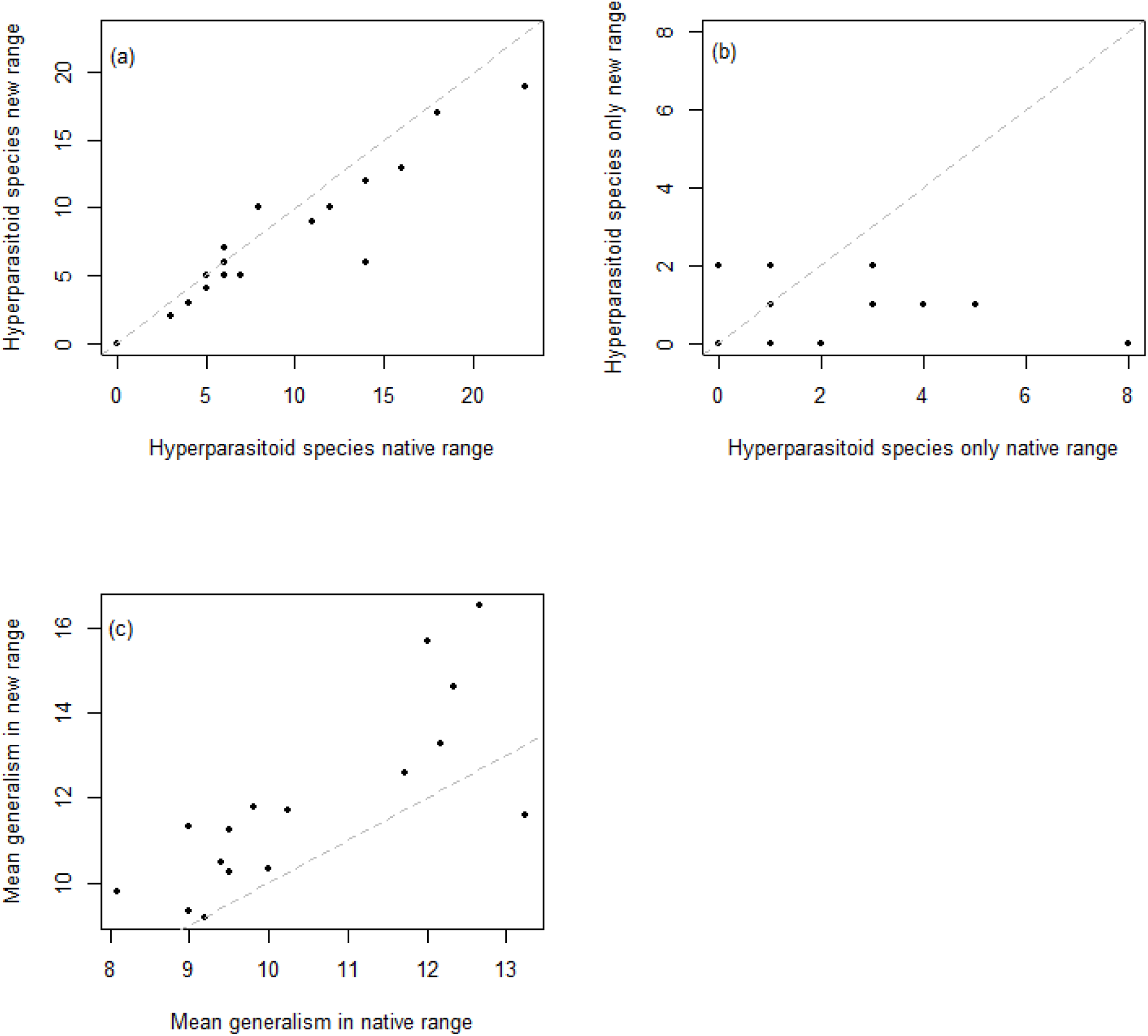
Number of hyperparasitoid species associated with the primary parasitoids introduced and established in continental North America to control aphids. (a) Total number of hyperparasitoid species in native and new ranges, and (b) the number of species which are found either only in the native range or only in the new range. (c) Mean number of hosts attacked per hyperparasitoid in the native and new ranges.

**Table 2.** Analyses and results. DER is the "degree of enemy release".

| Question | Value | Phylogenetic signal | P-value |
| --- | --- | --- | --- |
| Is there enemy release for established species? | PGLMM, $Z = 2.96$ | 0.23 | 0.003 |
| Is establishment affected by DER? | PGLMM, $Z = 0.82$ | 0.53 | 0.41 |
| Is establishment affected by the number of associations in the home range? | PGLMM, $Z = 0.24$ | 0.34 | 0.81 |
| Is control affected by DER? | PGLMM, $Z=0.84$ | 0.31 | 0.4 |
| Is there a correlation between the number of hyperparasitoids in the native and new range? | cor_phylo, 0.88 | 0.11 native, 0.11 new range | <0.0001 |
| Is there a correlation between the number of hyperparasitoids only in the native and only in the new range? | cor_phylo, -0.21 | 0.46 native, 0.08 new range | 0.43 |
| Does DER diminish through time? | cor_phylo, -0.06 | 0.13 | 0.81 |
| Does the proportion of mummy hyperparasitoids differ between the native and new ranges? | $t = 2.6$, $df = 13$ | 0 | 0.02 |
| Does the proportion of generalist hyperparasitoids differ between the native and new ranges? | $t = 4.3$, $df = 13$ | 0.42 | 0.0008 |

Whether the established species led to successful biological control was also not affected by the degree of enemy release (PGLMM, Z=0.84, *p*=0.4, with a phylogenetic random effect of 0.3).

The correlation between the number of hyperparasitoid species associated with the established primary parasitoids in the new range and in the native range was positive and strong (Pearson correlation was 0.88, *p* < 0.0001, with the phylogenetic signal of 0.11 for both native and new range) (Fig. 2a). This association is due to the fact that most associations between primary and hyperparasitoid species were the same in the native and new ranges, as evidenced by the weak association between the number of hyperparasitoids that are only found in the native range and the number only found in the new range (Pearson correlation of -0.21, *p* = 0.4, with a phylogenetic signal of 0.46 and 0.08 for the native and new range, respectively) (Fig. 2b). The year of parasitoid release was not related to the degree of enemy release for the established species (Pearson correlation was –3.02, *p* = 0.2) (Fig. S1). For the established parasitoid species, the proportion of hyperparasitoids that attack mummies rather than living aphids was greater in the new range than in the native range (t = 4.2, df = 13, *p* = 0.001). Finally, the hyperparasitoids in the new range included more generalists than in the native range (t = 4.3, df = 13, *p* = 0.0008) (Fig. 3c).

## Discussion

This study explored the enemy release hypothesis (ERH) for primary parasitoids of aphids that were introduced for biological control into North America. The data show that there was on average enemy release in the new range, but that this release did not affect establishment or successful biological control.

### Hyperparasitoid effects on primary parasitoids

ERH predicts that the chances for successful invasion, or establishment in the case of biological control agents, are increased if there are fewer natural enemies in the invaded range (Taylor 1955; Heimpel and Mills 2017). The 29 species of introduced aphid parasitoids studied here experienced different scenarios in the new range, with some encountering the same enemies as in the native range, some attacked by new species, and most experiencing varying degrees of enemy release (Fig. 2a, Table 1). However, there is no statistical evidence that establishment was affected by the degree of enemy release. This could indicate that the number of enemy species that a control agent encounters does not affect establishment, at least for the degree of enemy release found in this study. The relatively modest enemy release we found was largely due to the positive correlation between the degree to which hyperparasitoids were generalists and their geographical range (Fig. 1a); primary parasitoids that were attacked by many species of hyperparasitoids in their native range were attacked by many of the same hyperparasitoid species in the new range. The broad geographical ranges of hyperparasitoid generalists may be due to their having higher chances of natural or inadvertent human introductions and establishment, because they are more likely to find suitable hosts (“niche breadth–invasion success hypothesis”, Vazquez 2006). It is unlikely that hyperparasitoids were introduced together with primary parasitoids as part of biological control programs at least in recent years, because introduced biological control agents are strictly screened for the presence of hyperparasitoids (Heimpel and Mills 2017; Van Driesche et al. 2020). Nonetheless, early biological control programs might have been responsible for some hyperparasitoid introductions (Tougeron and Tena 2019), and inadvertent introductions are also possible when plant material is shipped or by other means (Murray and Mansfield 2015). It is also possible that establishment and control were not affected by the degree of enemy release because hyperparasitoids have positive effects on biological control by stabilizing the population dynamics of the control agent (Beddington and Hammond 1977; Rosenheim 1998; Heimpel and Mills 2017).

Records of hyperparasitism on biological control agents are incomplete, and we only had information on the possible presence of hyperparasitoid species rather than on their attack rates on control agents. Therefore, it is possible that we did not detect an effect of enemy release on the establishment and success of biological control agents because the data are incomplete (Tougeron and Tena 2019). Some groups of hyperparasitoids are extremely understudied, and in our literature search new host records for hyperparasitoids were mostly absent for recent years, suggesting a decrease in data collection on hyperparasitoid host ranges. There could also be bias in the literature; for example, more studies might be conducted in the new range following the introduction of a control agent than in the control agents native range. We tried to account for bias in the literature using the number of articles published and the year the species was described in our analyses. Nonetheless, limited information about hyperparasitoids might limit our ability to detect effects they have.

### Vulnerability to parasitism

In their analysis of introduced herbivores and their primary parasitoids, Cornell and Hawkins (1993) showed that the number of native parasitoids attacking invasive herbivores in a new range was positively correlated to the number in the native range; they referred to this as ’vulnerability to parasitism’. Superficially, we found a similar result for the higher trophic level of primary parasitoids and their hyperparasitoids (Fig. 2a). However, we show that correlation between the number of hyperparasitoids in the native and new ranges is the result of the same hyperparasitoid species occurring in both ranges; when asking this question for hyperparasitoids that only occur in the native or new ranges, the correlation disappears (Fig. 2b). Therefore, the correlation that we found between hyperparasitism in native and new ranges cannot be explained by vulnerability to parasitism as defined by Cornell and Hawkins (1993).

### Traits of enemies in the new versus native range

Introduced parasitoids did not only experience partial enemy release, the enemies they encountered differed in life-history traits. Higher proportions of mummy (vs. larval) hyperparasitoids were found in the new range, and since all of the hyperparasitoids that attack live aphids in our database are from the subfamily Charipinae, it means that enemy release was mostly due to reductions of this group. A similar finding was reported for the species *Lysiphlebus testaceipes* which invaded Benin (Hofsvang et al. 2014). Brodeur (2000) also made the observation that Charipinae are well represented in the native range of aphid primary parasitoid complexes and are rare in introduced complexes, and he called for comparative studies of hyperparasitoids in native and new ranges, as we have done. The relatively large number of mummy hyperparasitoids in the new range may also occur because these hyperparasitoids are more generalists (have greater numbers of hosts) (Fig 2c), as predicted by ERH (Keane and Crawley 2002). In addition to being mummy parasitoids, most of the hyperparasitoids in the new range (excluding two *Syrphophagus* species) were idiobiont ectoparasitoids, which further allows them to attack new species of hosts because their physiology does not need to be as tightly adapted to the host physiology (Askew and Shaw 1986; Brodeur 2000).

Our finding that introduced aphid parasitoids are attacked by numerous generalist hyperparasitoids suggests that biological control programs could have nontarget effects on the insect community, in particular other primary parasitoids through apparent competition (Van Nouhuys and Hanski 2000; Morris et al. 2001). Specifically, the control agents could increase the abundance of hyperparasitoids that then attack native parasitoids. The large number of generalist hyperparasitoids might also affect the evolution of the control agent, because generalist natural enemies change the selective pressure for traits that protect against attack (Joshi et al. 2010; Roy et al. 2011).

### Hyperparasitism rates and dynamics

Our data on the association between primary parasitoids and hyperparasitoids consists only of observed host records of the hyperparasitoid, and we do not have data on hyperparasitism rates. Although hyperparasitism rates are sometimes reported in field studies, they are usually reported only for the genus level of the primary parasitoid; species-level associations are hard to determine without molecular analyses to identify the species of parasitoid from which a hyperparasitoid emerged (Poelman et al. 2022). Therefore, a limitation of our study is the absence of hyperparasitism rates which could be used to determine the strengths of association between primary parasitoids and hyperparasitoids. The data that we could find show that hyperparasitism in the new range can be significant and similar to the native range. For example, *Trioxys pallidus* was introduced from Europe, where 2,230 mummies were collected in five different countries and more than 70% were hyperparasitized (Messing and Aliniazee 1989). In Oregon, USA, where this species was released to control the filbert aphid, *Myzocallis coryli*, over 60% hyperparasitism was observed in the first year after release. Despite high hyperparasitism, this species established and provides successful control (Hougardy and Mills 2009). Mackauer (1971) gives an example of the parasitoid *Aphidius smithi* which was released into Nova Scotia, Canada, to control pea aphids and did not establish, perhaps due to high pressure from hyperparasitoids, although evidence of high hyperparasitism was not shown and the author gives alternative explanations such as climate. High hyperparasitism rates did not explain the failure of biological control of the melon aphid *Aphis gossypii* in Hawaii. Results from a comparative study of the susceptibility to hyperparasitism of two parasitoid species introduced into Hawaii show that the more abundant species in the field is also more susceptible, making hyperparasitism an unlikely cause for the low abundance of the species that is less susceptible (Acebes and Messing 2013).

From the data available, genus-level hyperparasitism in the field is similar for the biological control agents in their native and new ranges (Angalet and Fuester 1977; Jacobson 2011). Furthermore, levels of hyperparasitism on biological control agents vary among locations, seasons, and years (Bosch et al. 1979; Zuparko 1995; Acheampong et al. 2012), as is found for parasitoids in their native ranges (Höller et al. 1994; Pons et al. 2011). Schooler et al. (2011) suggest that small populations of parasitoids, like those introduced for biological control, are especially susceptible to hyperparasitism. This suggests the strategy of releasing biological control agents to coincide with periods of low hyperparasitism. However, this would require more information than is presently available for many control agents and their hyperparasitoids.

## Conclusion

Our results show that enemy release may be present for introduced species of parasitoid biological control agents. Not only might there be fewer hyperparasitoid species, but their life history traits may differ, with more generalist hyperparasitoids in the new range. The increase in generalism we documented, however, was mainly due to a loss of specialist species in the new range and the presence of generalist species in both native and new ranges. Although there was some enemy release, it had no detectable effect on the rates of establishment or the success of biological control agents, as shown for other invasive species groups (Heger and Jeschke 2018). Our results, however, are for the presence/absence of hyperparasitoid species, rather than hyperparasitism rates, and more research is needed to better understand hyperparasitism rates in native and new ranges. More information on hyperparasitism rates is also needed to understand the community-level consequences of biological control that act by changing hyperparasitism on native species, such as apparent competition (Reynolds 1988; Tompkins et al. 2003).

## Acknowledgment

This work was funded in part by a Steenbock Professorship at the University of Wisconsin-Madison.

