## Supplementary figure 1 for "Enemy release of introduced parasitoids does not affect their establishment or success"

**Supplementary material**

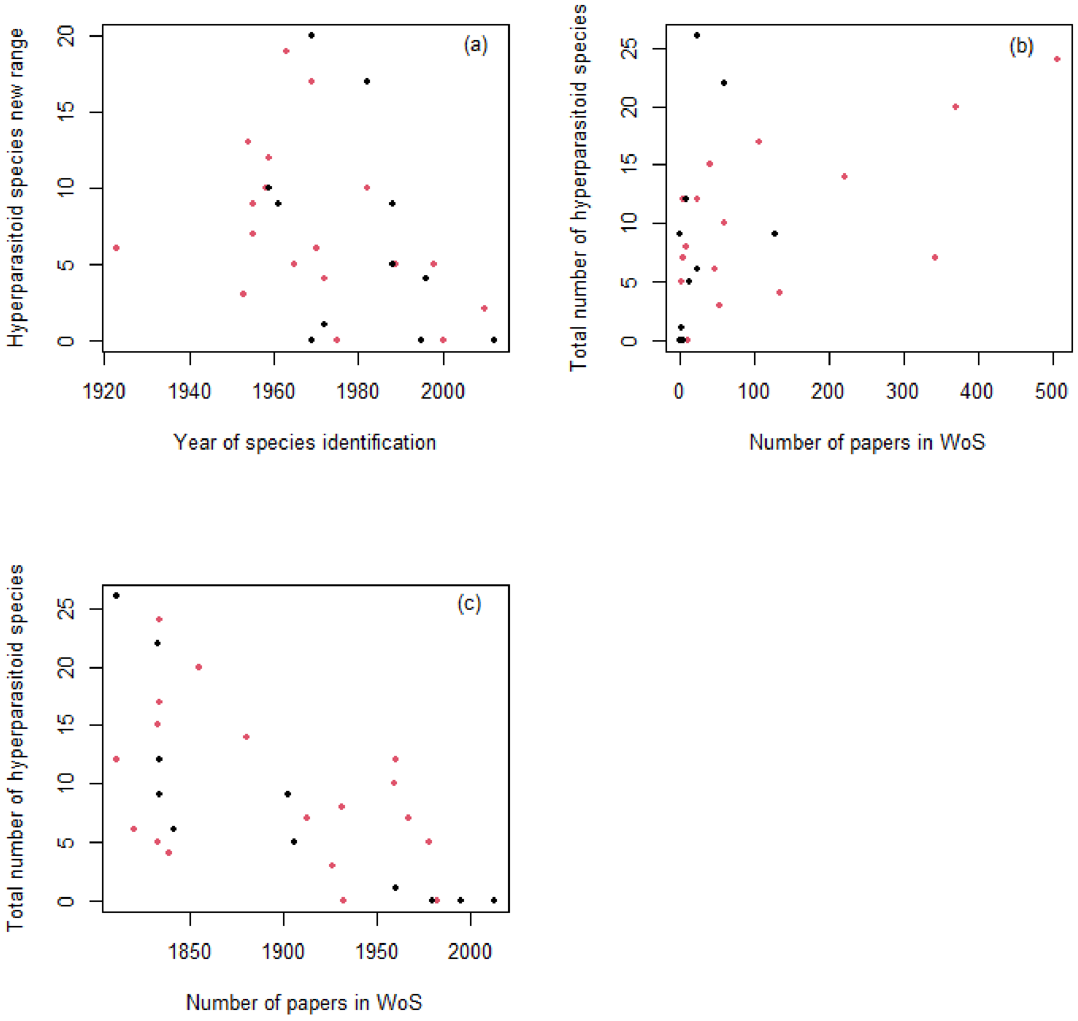


**Figure S1.** Number of hyperparasitoid species associated with the primary parasitoids introduced and established in continental North America to control aphids; species that did not establish are shown in black, and species which established are shown in red. (a) Number of hyperparasitoid species of primary parasitoids versus the year in which the primary parasitoid was introduced. (b) Total number of hyperparasitoid species versus the number of articles found in the Web of Science with the full name of the parasitoid (searched in February 2024). (c) Total number of hyperparasitoid species versus the year the primary parasitoid was first discovered.
